# Cell-type resolved prenylomics in mouse liver enabled by a low-input proteomics workflow

**DOI:** 10.64898/2026.10.23.761417

**Authors:** Harinarayanan Kottala, LeeAnn Higgins, Gavin Fredrickson, Mara T. Mashek, Douglas Mashek, Mark D. Distefano

## Abstract

Protein prenylation is a lipid modification crucial for membrane association and cellular signaling, but its dysregulation is linked to cancer and neurodegenerative diseases. Typical “prenylomic” workflows rely on metabolic labeling with bio-orthogonal isoprenoid probes followed by streptavidin enrichment; however, they generally require milligram-scale protein inputs (15–25 million cells), preventing the study of scarce primary or sorted cell populations. To overcome this limitation, we developed a highly sensitive, low-input prenylomic workflow combining optimized streptavidin enrichment with library-free data-independent acquisition (DIA) mass spectrometry. This method enables deep prenylomic coverage from just 50 µg of total protein input, a 20-fold reduction compared to prior methods. In COS7 cells, this workflow identified 91 prenylated protein groups, outperforming TMT-DDA methods that yielded 81 groups from 2 mg of input, using the same bio-orthogonal probe for enrichment, while coefficient of variation (CV) analysis demonstrated superior quantitative precision and reproducibility. Applied *in vivo* in mice following subcutaneous probe administration, the low-input methodology overcame sample input limitations to profile cell-type–specific prenylomes from dissociated liver tissue, identifying 29 enriched prenylated proteins in hepatocytes and 18 in macrophages. Pathway enrichment analysis showed that the identified proteins are consistent with the biology of the respective cell-types. This robust methodology significantly lowers the sample threshold for lipid proteomics, expanding the accessibility of prenylomics to limited primary cells and sorted tissue populations to study cell-type–specific prenylation dynamics in disease contexts.

## Introduction

Protein prenylation is a post-translational modification that attaches isoprenoid to the C-terminus of proteins containing a specific sequence.^1,2,3^ The lipids include either 15 carbon farnesyl or 20 carbon geranylgeranyl groups that become linked to specific cysteine residues via thioether bonds.^4,5^ The canonical CaaX box motif, where “C” is cysteine, “a” represents aliphatic amino acids, and X can be a variety of residues that direct either farnesylation catalyzed by farnesyltransferase (FTase) or geranylgeranylation catalyzed by geranylgeranyltransferase type I (GGTase I).^1,3^ Similarly, motifs including CC, CXC, or CCXX target proteins specifically for type II geranylgeranylation, a mechanistically more complex process mediated by geranylgeranyltransferase type II (GGTase II)^6^. A new GGTase complex has been reported recently, known as GGTase III, that has an orphan prenyl transferase alpha-subunit PTARI and the catalytic beta-subunit of GGTase II. GGTase III can specifically recognise the protein FBXL2. In general, prenylation promotes membrane association and mediates critical protein protein interactions.^1,2,3^

Inhibiting prenylation has been of long-standing interest in clinical studies for a number of diseases. Inhibition of farnesylation using farnesyltransferase inhibitors (FTIs) was initially the main focus of those studies.^7^ A Phase 2 clinical trial using the FTI Tipifarnib for H Ras driven head and neck cancer is currently in progress.^8^ Recently, the use of the FTI Lonafarnib was approved by the FDA for the treatment of Hutchinson-Gilford progeria syndrome, a premature aging disease.^9^ Overall, these studies have made it clear that a better understanding of which proteins are actually prenylated and how they vary in disease is required to develop more effective therapeutic strategies.^10^

Earlier methods to visualize prenylated proteins relied on autoradiographic techniques using tritiated isoprenoid precursors,^11,12,13^ as well as immunological approaches using isoprenoid directed antibodies.^14,15^ However, these traditional techniques suffered from significant limitations, including low sensitivity, long exposure times, high cost and irreproducibility. Mass spectrometry based chemical proteomics has since emerged as a powerful tool to overcome those limitations and profile and quantify prenylated proteins. Because direct identification of prenylated peptides remains difficult, and because most prenylated proteins contain well defined terminal sequences where the modification occurs, efforts typically focus on enriching intact prenylated proteins using bioorthogonal probes and comparing probe treated samples versus untreated controls rather than searching for modified peptides directly. For metabolic labeling, bioorthogonal probes bearing azide or alkyne groups have been extensively used in previous studies to enrich prenylated proteins. Following cellular incorporation, click chemistry conjugation to biotin or fluorophore handles enables selective enrichment and visualization.^16,17,18,19^ To enable relative quantification across conditions, early chemical proteomic approaches utilized probe enrichment followed by Stable Isotope Labeling by Amino Acids in Cell Culture (SILAC)^20^ or multiplexed isobaric tagging,^21,22^ using tandem mass tag-based (TMT) reagents. Those strategies employed alkyne-containing isoprenoid analogues paired with a biotin azide reporter and streptavidin pulldown.^23,24,25^ Using this approach, our group has successfully mapped and quantified the prenylome across diverse biological contexts, including COS7 cells, breast cancer models, brain derived cell lines, primary cells,^25^ and animal tissue models as well as investigating disease dynamics in Alzheimer’s disease,^26^ amyotrophic lateral sclerosis (ALS)^27^, and cancer.^28^ However, all previous prenylomic workflows suffer from high material requirements, typically demanding at least 1 to 2 mg of total protein input per sample.^25^ This requirement creates a major bottleneck when investigating biological material where input is limited including primary cells or sorted cells.

In this work, we describe a highly sensitive, low input chemical proteomic workflow utilizing an alkyne-containing isoprenoid probe paired with scaled down streptavidin magnetic bead enrichment and Data Independent Acquisition Liquid Chromatography Mass Spectrometry (DIA LC-MS2) processed via DIA-NN.^29^ To develop this new strategy, we first, evaluated the analytical depth, quantitative precision, and reproducibility of this workflow in COS7 cells pretreated with Lovastatin^25^ using a greatly reduced protein input of 50 µg, directly benchmarking its performance against established TMT DDA methods. We then applied this high sensitivity pipeline in vivo to characterize the prenylomes of two different cell types in mouse livers including hepatocytes and macrophages. Metabolic labeling efficiency was initially evaluated in those cells via in gel fluorescence with a TAMRA azide reporter tag. Next, quantitative profiling of the prenylomes was carried out via our new DIA LC-MS2 method. Finally, bioinformatic analysis was performed to reveal correlations between the prenylated proteins identified and the biological functions of the hepatocytes and macrophages studied.

## Results and Discussion

### Comparing DIA LC-MS2 and TMT-SPS-MS3 workflows at 2 mg inputs

To evaluate data-independent acquisition (DIA) paired with DIA-NN data processing for prenylomic profiling, we first tested performance under standard input conditions (2 mg total protein). COS7 cells pretreated with lovastatin were metabolically labeled with either the C15AlkOPP probe or the natural substrate farnesyl diphosphate (FPP) as a control, followed by biotinylation and streptavidin enrichment. To enable a direct comparison, an aliquot from the same sample was used for TMT labeling and SPS-MS3 analysis employing our previously established method,^25^ while the remainder was processed via the label-free DIA LC-MS2 workflow (**Figure 1A**). This initial test served to verify that DIA acquisition and DIA-NN processing can be effectively employed in the context of prenylomic analysis before implementing a lower input.

**Figure 1:**
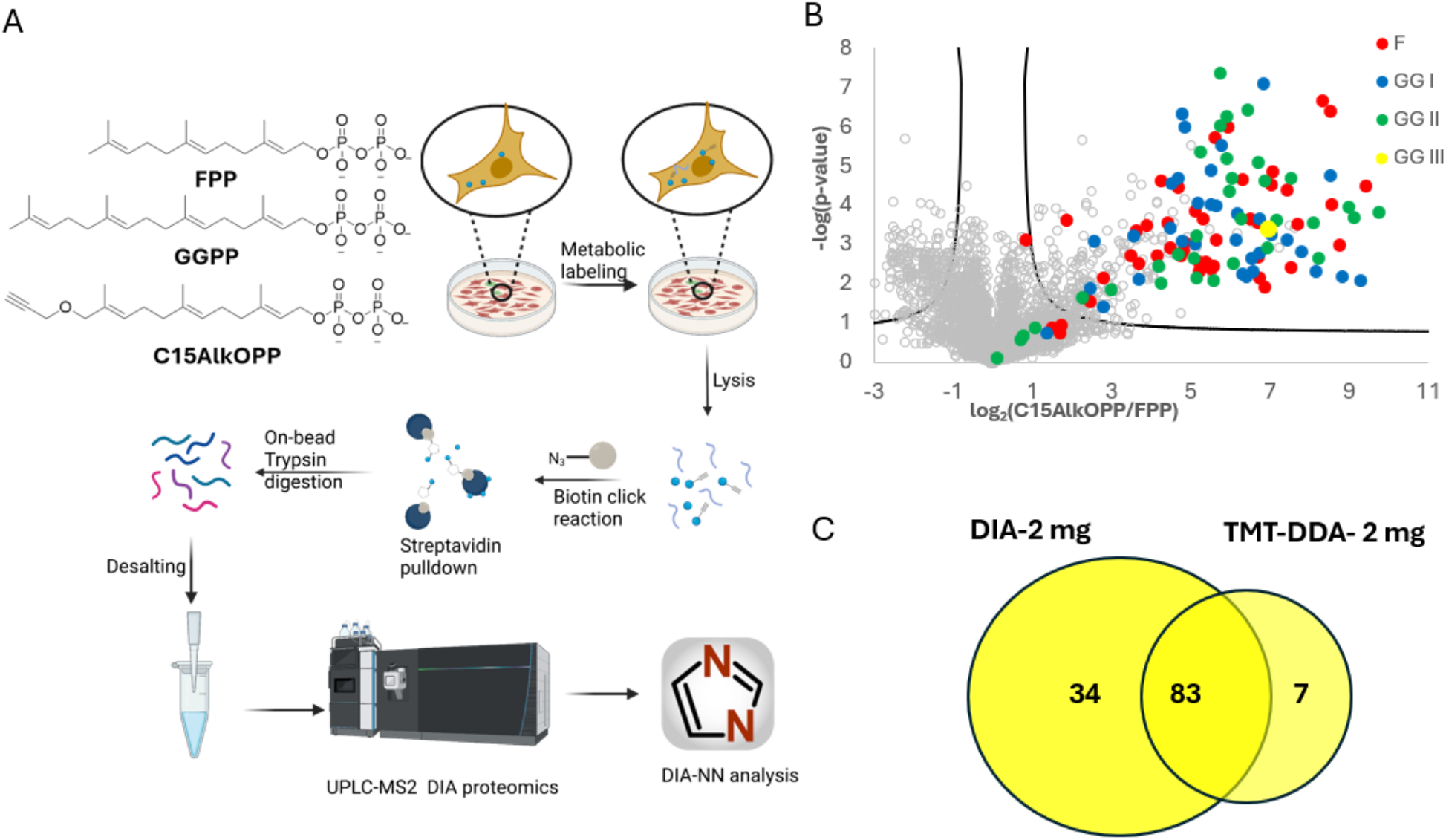
**A)** Schematic representation of the prenylomic workflow utilizing bio-orthogonal isoprenoid probe (C15AlkOPP) for metabolic labeling, click chemistry-mediated biotinylation, streptavidin enrichment, on-bead digestion, and DIA LC-MS2 analysis processed via DIA-NN. **B)** Volcano plot showing differential enrichment of prenylated proteins from COS7 cells pretreated with statin (lovastatin) and labeled with the C15AlkOPP probe relative to FPP controls using 2 mg total protein input with DIA mass spectrometry and DIA-NN processing. Enriched targets are color-coded by prenylation substrate class: FTase (F, red), GGTase I (GG I, blue), and GGTase II (GG II, green). **C)** Venn diagram illustrating the overlap in identified prenylated protein groups between DIA-MS2 (2 mg input) processed via DIA-NN and the established TMT-SPS-MS3 method (2 mg input) processed using MaxQuant.

A volcano plot of the C15AlkOPP-labeled sample processed via DIA-NN showed significant enrichment of 117 known prenylated proteins (ungrouped) relative to FPP controls, spanning farnesylated (F), type I geranylgeranylated (GG I), and type II geranylgeranylated (GG II) substrates (**Figure 1B**). When comparing ungrouped prenylated proteins via Venn diagrams with the parallel sample aliquot analyzed by TMT-SPS-MS3 (which identified 92 proteins), 83 proteins were shared between both methods (**Figure 1C**). Overall, these results demonstrate that DIA mass spectrometry paired with DIA-NN processing yields coverage superior to the established TMT-SPS-MS3 method under standard input levels, supporting its implementation in a low-input workflow.

### Low-input profiling of prenylated proteins using DIA analysis

To evaluate the performance of the new workflow using lower protein input, the prenylome of COS7 cells treated with the C15AlkOPP probe following lovastatin pretreatment was studied. In-gel fluorescence analysis showed that C15AlkOPP exhibited superior labeling efficiency in the presence of lovastatin (**Figure 2A**). The goal of these experiments was to substantially lower the protein input down to 50 µg, aiming for a 20-fold reduction in sample amount compared to previously published analyses, without sacrificing proteomic depth or quantitative fidelity. Downstream sample processing was performed as outlined in **Figure 1A**. The enrichment volume was scaled down to 100 µL, and to facilitate small sample size, streptavidin magnetic beads were used for enrichment. DIA LC-MS2 was performed using an Orbitrap Eclipse instrument, and the data was analyzed using DIA-NN (v 2.2.0). The data employed for comparison was taken from a previously published paper from our group with the same cell-line and treatment.^25^ Directly benchmarking novel chemical proteomic pipelines against previous datasets is inherently complex due to variations in instrumentation generations, data acquisition strategies, and downstream identification and quantification algorithms. Nevertheless, a general comparison shows substantial agreement in prenylated protein identification. While our previously published prenylomic method which used a Tandem Mass Tag Data-Dependent Acquisition (TMT-DDA) workflow identified 81 prenylated protein groups from 2 mg of protein input, the new workflow identified and quantified 91 prenylated protein groups from 50 µg of starting protein input (**Figure 2B**). The overlap between these two acquisition methods is illustrated in a Venn diagram of the ungrouped proteins (Figure 2C). The 81% overlap between the two methodologies shows strong concordance between the two approaches despite the much lower sample input in the new workflow. A coefficient of variation (%CV) analysis was conducted to measure quantitative fidelity under the decreased sample input. A box plot comparison of %CV values (**Figure 2D**) revealed a lower median %CV for the 50 µg workflow compared to the previously published 2 mg workflow, demonstrating that the combination of scaled-down magnetic bead enrichment and DIA-NN processing delivers improved quantitative precision.

**Figure 2:**
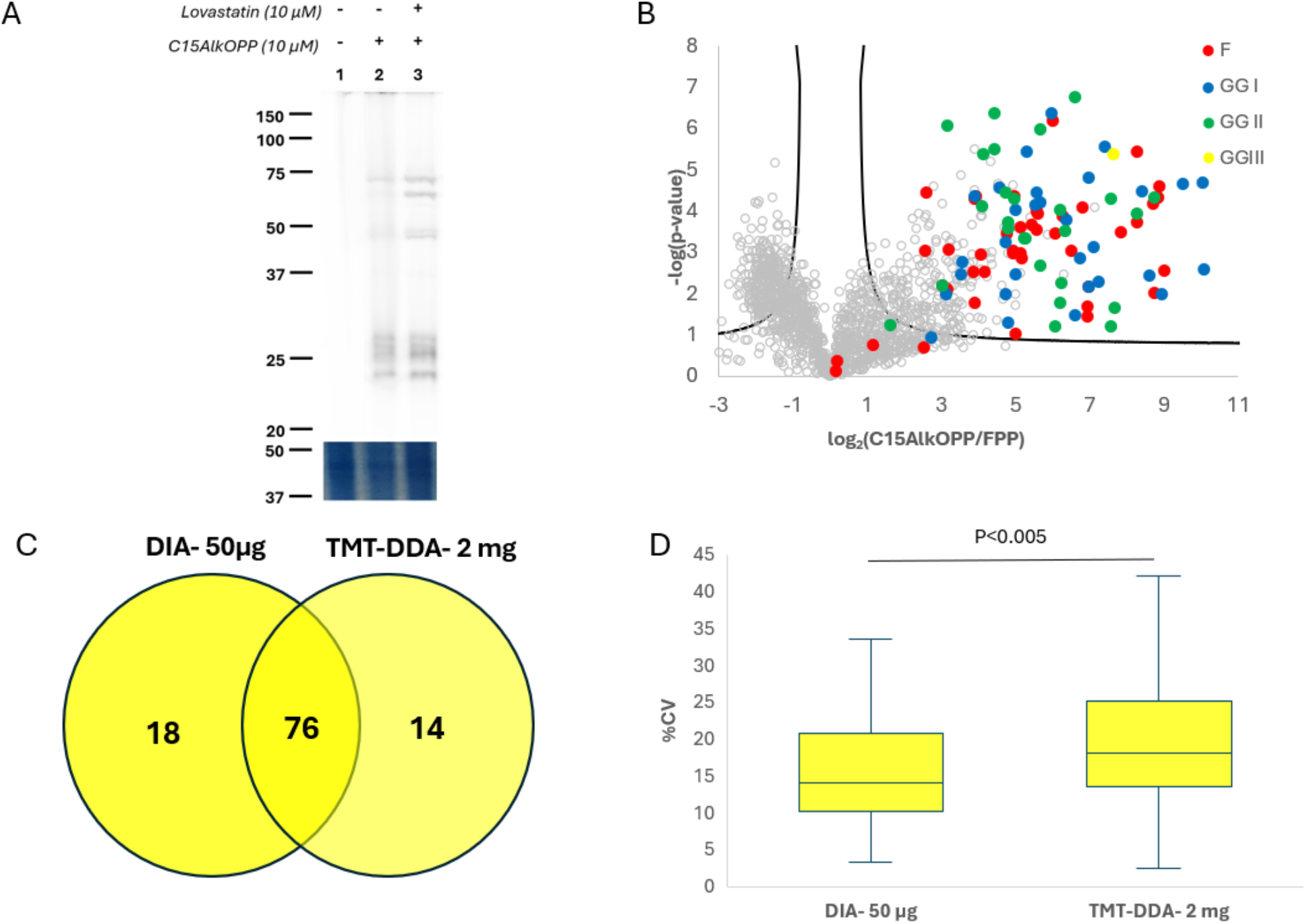
**A)** In-gel fluorescence analysis (top) depicting metabolic incorporation of bio-orthogonal isoprenoid probes (C15AlkOPP) in COS7 cell lysates with or without Lovastatin pretreatment, paired with Coomassie blue staining (bottom). **B)** Volcano plot of prenylomic profiling in COS7 cells showing relative enrichment in C15AlkOPP-treated samples using 50 µg sample input. Enriched targets are categorized by prenylation class: farnesylated (F, red), geranylgeranylated type I (GG I, blue), and geranylgeranylated type II (GG II, green) (FDR= 1%, s0=0.5). **C)** Venn diagram illustrating the overlap of ungrouped prenylated protein identifications between the scaled-down DIA workflow using 50 µg protein input and the traditional TMT-DDA workflow using 2 mg protein input. **D)** Box plot comparison of quantitative precision measured by percent coefficient of variation (%CV) across replicates for the 50 µg DIA workflow versus the 2 mg TMT-DDA workflow (*P* < 0.005). The horizontal line indicates the median.

### Cell-type specific prenylomics of mouse liver

Next, the ability of the new workflow to capture cell-type–specific prenylomes *in vivo* using primary cell populations isolated from mouse liver was evaluated (**Figure 3A**). Importantly, while hepatocytes yield milligram amounts of protein per mouse, isolation of primary intrahepatic macrophages typically yields only 400,000–700,000 cells per liver, producing 15–20 µg of total protein. The ability to perform prenylomic analysis using much lower amounts of protein from small cell populations was one of the major motivations for the development of the new workflow presented here.

**Figure 3:**
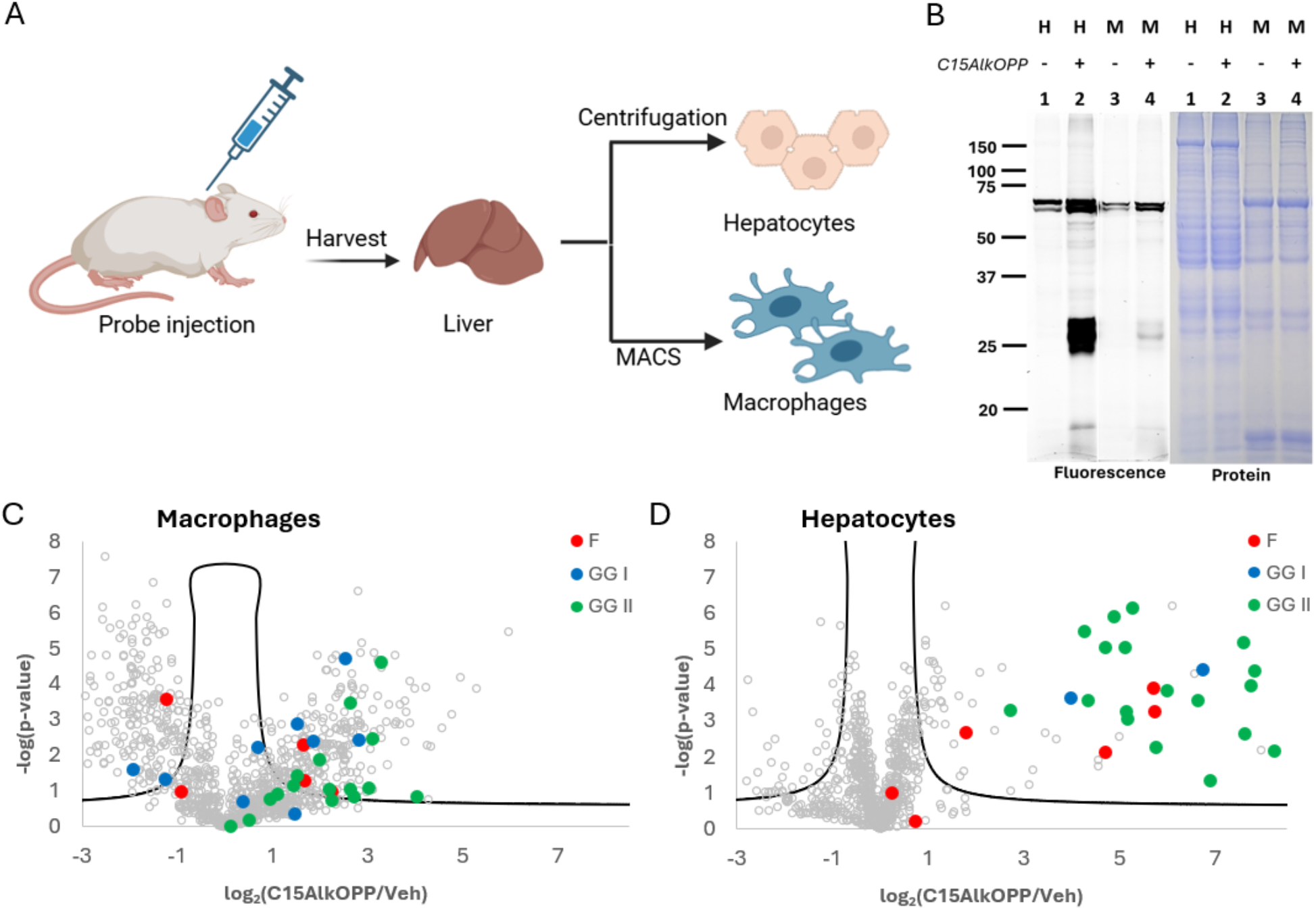
**A)** Schematic representation of the in vivo experimental pipeline showing subcutaneous C15AlkOPP probe administration in mice, liver tissue harvest, and isolation of primary hepatocytes via centrifugation and resident macrophages via magnetic-activated cell sorting (MACS). **B)** In-gel fluorescence visualization (left) following TAMRA-azide click chemistry conjugation, alongside total protein loading control (right), comparing metabolic probe labeling efficiency in isolated hepatocytes (H) and macrophages (M). **C)** Volcano plot of differential prenylated protein enrichment in primary resident macrophages, highlighting identified farnesylated (F, red), type I geranylgeranylated (GG I, blue), and type II geranylgeranylated (GG II, green) targets (FDR= 5%, s0=0.5). **D)** Volcano plot of differential prenylated protein enrichment in isolated hepatocytes, color-coded by target prenylation class (FDR= 5%, s0=0.5).

Prior to mass spectrometry, metabolic labeling efficiency was evaluated via in-gel fluorescence using click chemistry and a TAMRA-azide reporter (**Figure 3B**). Lysates from hepatocytes obtained via in vivo metabolic labeling with C15AlkOPP-exhibited robust labeling dominated by several bands near ~25 kDa, consistent with the molecular weight of small GTPases (e.g., Ras, Rab, and Rho families). Macrophage labeling revealed a similar pattern including bands in the 25 kDa region although the overall fluorescence intensity across all molecular weight ranges was much weaker; vehicle-treated controls showed negligible signal in the 25 kDa region. It should be noted that each lane in the gel contained 25 µg (based on BCA assay) and Coomassie blue staining suggested comparable protein loading in all lanes (Figure 3B). The stronger labeling intensity in hepatocytes relative to macrophages likely reflects their higher metabolic activity and probe uptake dynamics *in vivo*.

Next, the prenylated proteins from each cell type were profiled using the new workflow. For this analysis, lysates were used from cells pooled from three mice. Following the click reaction and enrichment steps, the resulting samples were divided into three technical replicates prior to desalting, and DIA LC-MS2 analysis was performed as described above. In hepatocytes, the analysis identified 29 significantly enriched prenylated proteins in probe-treated samples (**Figure 3D**). These targets include representatives from all three prenylation classes, dominated by type II geranylgeranylated (GG II / Rab family) proteins. In primary resident macrophages, 18 prenylated proteins exceeded the significance threshold, also spanning farnesylated (F), geranylgeranylated type I (GG I), and GG II classes (Figure 3C). This lower target count and reduced fold-change span directly mirror the weaker fluorescence signals observed by in-gel analysis. Because primary macrophages yield only 15–20 µg of protein per mouse, successfully mapping their prenylome demonstrates the practical utility of this low-input workflow. Comparative analysis with a Venn diagram showed that 14 prenylated proteins were shared between both cell populations, while 15 were unique to hepatocytes and 4 were unique to macrophages (**Figure 4C**). While this initial analysis demonstrates robust profiling from limited input, expanding this workflow to additional biological replicates in future studies will be essential to evaluate animal to animal variability.

**Figure 4:**
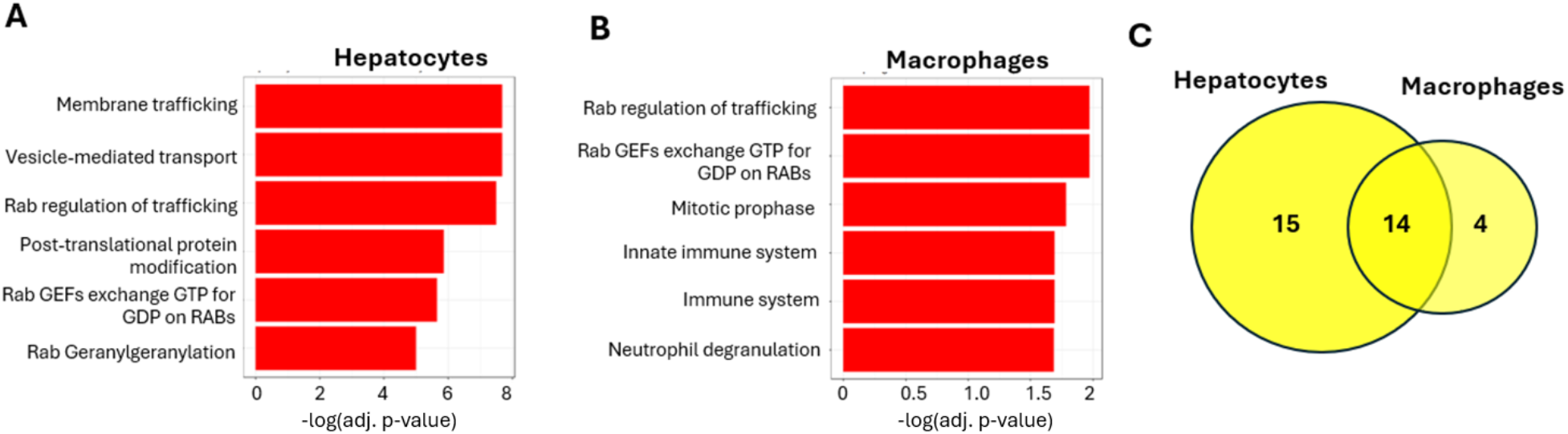
**A)** Bar chart displaying the top 6 significantly enriched biological pathways identified in hepatocytes, ranked by −log_10_(adjusted p-value). B) Bar chart displaying the top 6 significantly enriched pathways identified in macrophages, ranked by −log_10_(adjusted p-value). C) Venn diagram showing shared (14) and unique ungrouped prenylated targets identified between isolated hepatocytes (15 unique, ungrouped) and macrophages (4 unique, ungrouped).

To explore the biological significance of the prenylomic results, Reactome pathway enrichment analysis was performed. The analysis of the 29 prenylated targets in hepatocytes revealed the top 6 most significant pathways which were heavily associated with cellular transport activity (**Figure 4A**). These top-ranked terms included “Membrane trafficking”, “Vesicle-mediated transport”, and “Rab regulation of trafficking”, as well as “Post-translational protein modification” and “Rab Geranylgeranylation”. However, the top 6 enriched pathways for the 18 macrophage targets reflected both core trafficking machinery functions as well as immune-specific processes (**Figure 4B**). In particular, while core vesicular transport terms (“Rab regulation of trafficking and Rab GEFs exchange GTP for GDP on RAB proteins) were highly ranked in both cell types, the macrophage prenylome showed preferential top-ranking for immune functional signatures, including “Innate immune system”, “Immune system”, and “Neutrophil degranulation”.

Collectively, these bioinformatic data demonstrate that even with low protein inputs isolated from primary macrophages, the workflow successfully captures cell-type-appropriate pathways within the liver microenvironment.

## Materials and Methods

### Metabolic labeling in cultured cells

COS7 cells were seeded at a density of 0.9–1 × 10^6^ cells in 100-mm dishes containing 10 mL of DMEM supplemented with 10% fetal bovine serum (FBS) and 1% penicillin–streptomycin 24 hours prior to treatment. For treatment, the media was replaced with 5 mL of fresh media, and cells were pre-treated with 10 μM lovastatin (Cayman chemical, #10010338) for 6 h (37 °C, 5% CO_2_). Following pre-treatment, C15AlkOPP or FPP was added to the media to a final concentration of 10 µM. Finally, after 24 hours of metabolic labeling, the media was removed, and the cells were washed twice with 1 mL of PBS for each plate and harvested by scraping them off the plates. Cell pellets were stored at −80 °C until further use.

### In-vivo labeling in Mouse and isolation of hepatocytes and macrophages

A total of 6 male mice at 10-12 weeks of age were used. For each condition, three mice were injected subcutaneously, in the interscapular region, with 150 μL of sterile-filtered C15AlkOPP (30 mM with 4.8 mM NH_4_HCO_3_ in PBS), and three mice were injected with 150 μL of sterile-filtered vehicle solution (4.8 mM NH_4_HCO_3_ in PBS). The treatment was terminated 24 h after a single injection or 72 h after three injections administered every 24 h. Hepatocytes were isolated by the collagenase perfusion method.^30^ Briefly, mice were perfused with collagenase through the hepatic portal vein, and livers were collected and minced. Subsequently, non-parenchymal cells were separated from hepatocytes by centrifugation at 50 × g, and the supernatant containing non-parenchymal cells was used for macrophage enrichment. Non-parenchymal cells were washed with PBS containing 0.5% BSA and 2 mM EDTA and incubated with anti-F4/80 microbeads (Miltenyi Biotec, #130-110-443) for 15 min. Cells were washed and loaded onto LS columns in magnetic MACS separators (Miltenyi Biotec). After three washes, purified macrophages were collected.

### In-gel fluorescence analysis

In-gel fluorescence analysis was performed according to previously established protocols.^25^ Cell pellets were resuspended in lysis buffer (200 units/nL benzonase nuclease, 1X protease inhibitor cocktail, and 1% SDS in PBS buffer) and lysed on ice via sonication (6–7 pulses, 3 s on, 7 s off). Following protein quantification by BCA assay, 50 µg of protein was subjected to a click reaction for 1 h at room temperature using 25 µM TAMRA-PEG3-N_3,_ (Broadpharm, # BP-22479) 1 mM TCEP, 0.1 mM TBTA, and 1 mM CuSO_4_. Proteins were then precipitated using a ProteoExtract kit (Calbiochem, # 539180) and resuspended in SDS loading buffer (50 mM Tris-HCl pH 6.8, 2% SDS, 10% glycerol, 0.02% bromophenol blue). Samples were resolved on 12% SDS-PAGE gels, and TAMRA fluorescence was visualized with a Typhoon FLA 9500 scanner. Image analysis was conducted using ImageJ. Following fluorescence imaging, the gel was stained with Coomassie Brilliant Blue R-250 to visualize total protein (0.2% (w/v) Coomassie Brilliant Blue R-250, 10% (v/v) CH_3_CO_2_H, 45% (v/v) methanol, and 45% H_2_O). The gel was subsequently destained using destaining solution (10% (v/v) CH_3_CO_2_H, 30% (v/v) methanol, and 60% (v/v) H_2_O) until the protein bands were clearly visible.

### Enrichment of Prenylated Proteins

Cells were lysed as described in the previous section. For each sample, lysate containing 50 µg of protein in 25 µL was subjected to CuAAC with 100 µM biotin-N3 (Broadpharm, #BP-22119), 1 mM TCEP, 0.1 mM TBTA, and 1 mM CuSO_4_ for 90 min at room temperature. Following the reaction, proteins were precipitated using a ProteoExtract kit (Calbiochem, # 539180) and resuspended in 100 µL of PBS containing 1% SDS. Protein concentrations were again measured using a BCA assay.

Equal amounts of biotinylated protein (typically 50 µg) were adjusted to a final volume of 100 µL and incubated for 3 h with pre-washed Pierce Streptavidin Magnetic Beads (Thermo Scientific, #88816), using 100 µL of bead slurry per sample. The beads were washed three times with 400 µL of PBS containing 1% SDS, followed by one wash with PBS, three washes with 8 M urea in 50 mM TEAB, and three washes with 50 mM TEAB. On-bead digestion was carried out overnight at 37 °C with 0.4 µg sequencing-grade trypsin (Promega, #V5111) in 50 µL of 50 mM TEAB. Peptides were extracted from the beads by sequential treatment with 100 µL of H_2_O containing 0.1% HCO_2_H and 100 µL of 30% CH_3_CN in H_2_O.

The three solutions were combined, dried by lyophilization, and resuspended in 60 µL of 0.2% HCO_2_H in water. Peptides were desalted using StageTips containing two layers of SDB-RPS extraction disks (Empore; 47 mm diameter, 0.5 mm thickness, #66886-U). Peptide solutions were loaded onto the stage tips and centrifuged till all the solution passed through the membrane. The StageTips were then washed twice with 60 µL of 5% CH_3_CN /0.2% HCO_2_H in H_2_O, followed by 60 µL of CH_3_CN. Bound peptides were then eluted with 60 µL of 50% CH_3_CN containing 5% NH_4_OH in H_2_O and dried by lyophilization.

### Liquid Chromatography-Tandem Mass Spectrometry (LC-MS/MS)

Dried peptides were resuspended in H_2_O containing 2% CH_3_CN and 0.1% HCO_2_H at the desired concentration, and 75 ng of peptides were injected for LC-MS/MS analysis. Peptides were separated using an Ultimate 3000 RSLCnano UHPLC system (Thermo Scientific) equipped with a self-packed C18 analytical column (100 µm id × 40 cm) containing ReproSil-Pur C18-AQ resin (1.9 µm, 120 Å; Dr. Maisch). The column was maintained at 55 °C. Mobile phase A consisted of 0.1% HCO_2_H in H_2_O, and mobile phase B consisted of 0.1% HCO_2_H in CH_3_CN. The flow rate was 350 nL/min during loading and equilibration and 315 nL/min during the analytical gradient. Peptides were separated using a 5–35% B gradient over 64 min.

Data were acquired using an Orbitrap Eclipse Tribrid mass spectrometer (Thermo Scientific) in positive ion mode using DIA. MS1 scans were collected from *m/z* 390–990 at a resolution of 60,000, with an AGC target of 4 × 10^5^ and a maximum injection time of 50 ms. Fragmentation was performed by HCD at a normalized collision energy of 33%. DIA MS2 spectra were acquired using 74 variable isolation windows spanning the MS1 range at a resolution of 30,000. The MS2 AGC target was 4 × 10^5^ with a maximum injection time of 54 ms, resulting in a cycle time of approximately 3 s.

### Data Processing and Bioinformatics Analysis

Raw files were analyzed using DIA-NN (v 2.2.0) against the non-redundant Homo sapiens reference proteome from UniProt (UP000005640) for COS7 samples and *Mus musculus* reference proteome from UniProt (UP000000589) for mouse samples. An *in silico* spectral library was generated using deep-learning-based predictions. The search parameters were set to trypsin digestion with up to one missed cleavage, precursor lengths of 7–30 amino acids, precursor *m/z* values of 300–1800, charge states of 1–4, and fragment *m/z* values of 200–1800. N-terminal methionine excision and methionine oxidation were included as variable modifications.

The predicted library was used for peptide and protein quantification with match-between-runs enabled. Quantification was performed using the QuantUMS high-precision algorithm with retention time-dependent cross-run normalization. Precursor and protein-group FDRs were both controlled at 1%.

Statistical analysis was performed using Perseus (v 2.1.3.0). Protein intensities were log2-transformed, and protein groups with more than four missing values out of the eight measurements were removed. Remaining missing values were imputed from a normal distribution using a width of 0.3 and a downshift of 1.8. Differential enrichment of prenylated proteins was assessed using a two-sample *t*-test with a permutation-based FDR cutoff of 5%.

### Pathway Enrichment Analysis

Functional pathway enrichment analysis of significantly enriched prenylated proteins in isolated hepatocytes and primary resident macrophages was conducted in R using the enrichR package.^31,32,33^ Prior to analysis, protein identities were converted to official gene symbols, standardized to uppercase, and deduplicated. To control for over-representation bias inherent to prenylated proteins and ensure that enriched terms reflect cell-type-specific biology, background correction was implemented by supplying a comprehensive reference list of known prenylated proteins (Obtained by filtering all known prenylated proteins from UniProt) as the custom background universe in enrichR. Gene sets were queried against the Reactome database (Reactome_Pathways_2024). Pathways were considered statistically significant at a Benjamini-Hochberg adjusted *P*-value threshold of adjusted p-value < 0.05. For visualization, the top six significant terms per cell population were ranked by −log_10_(adjusted p-value) and plotted using ggplot2.

## Acknowledgements

The authors thank Department of Chemistry at University of Minnesota for the Doctoral Dissertation Fellowship. Center for Proteomics and Metabolomics (CMSP) at University of Minnesota is acknowledged for their support with the mass spectrometer used in the proteomic experiments.

